# Taxanes act as vascular disrupting agents and increase rate of metastasis when combined with anti-angiogenic therapy

**DOI:** 10.1101/2022.07.18.500307

**Authors:** Rajender Nandigama, Mathias Kallius, Katharina Hemmen, Shaoli Das, Jürgen Pinnecker, David Ascheid, Verena Burkhard, Hla Ali, Johannes Rainer, Daniela Scheld, Sabine Herterich, Alma Zernecke-Madsen, Olaf Penack, Stefan Diller, Kevin Camphausen, Süleyman Ergün, Uma Shankavaram, Katrin Heinze, Freddy E. Escorcia, Erik Henke

## Abstract

Taxanes are known to have a profound effect on endothelial cells and the vasculature even at low doses. Here, we show that taxanes, rather than being anti-angiogenic, function more as vascular disrupting agents (VDAs), although they exert a different mechanism of vascular permeabilization when compared to traditional VDAs such as combretastatins. In the tumor context, this VDA-effect leads to a rapid vascular collapse and acute hypoxia. Concomitant treatment with anti-VEGF drugs aggravates hypoxia by blocking vasculogenic rescue mechanisms. While this results in a strong growth-suppressing effect on the tumor, it also increases its invasiveness and metastatic potential. We demonstrate that combination of anti-angiogenic drugs with taxanes blocks tumor reperfusion, intensifies intravasation of circulating tumor cells (CTCs) and strongly increases metastasis. Anti-VEGF drugs are commonly applied in combination with cytotoxic drugs including taxanes. Our findings have significant implications for the clinical use of this drug combination.

## Introduction

Anti-angiogenic drugs continue to play an important role in the management of advanced malignancies. The most frequently applied anti-angiogenic drug, Avastin (Bevacizumab) a VEGF-A sequestering antibody, is commonly not used as a stand-alone therapy but in combination with cytotoxic drugs like paclitaxel in ovarian cancer or 5-fluorouracil-based regiments in colorectal cancer (1-5). However, it has been demonstrated that various cytotoxic chemotherapeutics have anti-angiogenic properties by themselves (6-8). In particular, microtubule-binding drugs, like taxanes, appear to have a significant effect on the tumor vasculature although the exact mechanisms remain elusive (reviewed in (9)). These drugs have the potential to rapidly reduce tumor perfusion, consequently blocking supply with nutrients and oxygen. Consistent with this phenomenon, significantly elevated levels of pro-angiogenic cytokines, and increased numbers of circulating endothelial (CEC) and endothelial progenitor cells (CEPs) have been reported in patients following chemotherapeutic treatment (10, 11). Both effects can be interpreted as a response of the tumor to acute hypoxic stress. CEPs are therein seen as a rescue mechanism, to patch the damaged vasculature in response to a spike in pro-angiogenic signaling molecules secreted by the tumor cells under acute hypoxia. If this rescue mechanism is blocked, e.g. by co-administration of anti-angiogenic drugs, hypoxia is further aggravated in the tumor (12). While this seems to improve response of tumors to therapy, it also might activate an alternative escape mechanism: increased invasiveness and intravasation, which again could lead to enhanced metastasis.

Although addition of anti-angiogenic drugs to taxane-based regiments shows often robustly improved response rates and results in significantly increased progression free survival (PFS) it only moderately benefits overall survival (OS) (3, 4). A possible explanation might be that addition of anti-angiogenic drugs initiates mechanisms that only later manifest in detrimental effects, negating the initial benefits of the co-treatment. Increases in metastatic seeding would be such a delayed negative effect. Moreover, as the combination of anti-angiogenics with taxanes is used in the management of metastatic disease a further increase in metastatic seeding might be difficult to detect in patients while they are still monitored for progression.

We explored the effects of taxanes on tumor blood vessels in the context of co-application of anti-angiogenic drugs. Our focus were consequences of this drugs alone or in combination on vascular integrity, tumor supply and activation of pro-metastastic mechanisms. In mice, paclitaxel treatment indeed causes a rapid irreversible collapse of the tumor vasculature. This is caused by a vascular disrupting effect taxanes trigger in activated blood vessels. Dependence of this VD effect on pro-angiogenic stimulation explains the observed selectivity for the tumor vasculature, as other organs were less affected. Co-administration of anti-angiogenic drugs blocked vasculogenic rescue mechanisms and resulted in a strong pro-metastatic molecular response, that also manifested in the release of tumor cell in the circulation (CTCs) and an increase in pleural metastases. Importantly, tumor expression data from clinical trials demonstrated that in patients, too, markers for hypoxia, invasiveness, metastasis and reduced survival increased under combination therapy with bevacizumab/taxanes.

## Material and Methods

If not otherwise indicated, chemicals were acquired from standard commercial suppliers (SigmaAldrich, Merck, Carl Roth). Paclitaxel was dissolved in 50% v/v cremophore EL in EtOH, stored at -20°C and diluted in 3 volumes of sterile PBS prior to intraperitoneal injection. The pro-drug combretastatin A4-phosphate (CA4P, Fosbretabulin) was purchased from Selleckchem (Houston, TX). CA4P was dissolved in sterile PBS, stored at - 20°C and applied by intraperitoneal injection at the given dosages.

### Tumor models and treatment

All experiments involving animals were reviewed and approved by the Institutional Animal Care and Use Committee at MSKCC. The experiments were performed in accordance with relevant guidelines and regulations.

*Tumor engraftment:* LLC (1×10^6^ cells in matrigel/PBS 1:1) tumors were generated by subcutaneous injection in the dorsal region of female C57Bl/6J mice. 4T1 (1×10^5^ cells in PBS) breast adenocarcinomas were generated by injection of cells into the inguinal mammary fat pad of female BALB/c mice. BALB/c and C57Bl/6J were acquired from Jackson Labs, Bar Harbor, ME.

All animals in the individual experiments were of the same age and sex. For each experiment tumor bearing mice were randomly assigned to the different treatment groups just prior to the start of treatment. In treatment studies where tumor growth was a critical outcome assessment of tumor size was performed blinded by a second researcher.

*Exclusion of data:* Animals that never developed tumors due to take rate lower than 100 % were excluded from the studies. All data from animals that died or had to be sacrificed for health reasons prior to the scheduled termination of the experiment was excluded.

#### Tumor treatment

*Dosing of PTX:* PTX Dosage in mice (20 mg/kg BW twice in 6 days) was adjusted from standard dosing schedules inpatients. Patients that undergo bevacizumab paclitaxel combination treatments regularly receive both drugs concomitantly at the same day (3-5). Paclitaxel is given in these combination regiments at doses of 90 – 200 mg per square meter of body-surface area (app. 2.3 – 5 mg/kg BW for a 75 kg /170 cm patient). The standard conversion factor of drug dose from human to mouse recommended also by the NIH is 12.3 (13). We therefore used a dose of 20 mg/kg BW twice in 6 days, i.e. a lower individual dose repeated more frequently, to adjust for the faster growth of tumors in the mice but to still remain in the overall dosing range applied to human patients.

VEGF-A antibody mG6-31 (Genentech, San Francisco, CA) was applied by i.p. injections at 6 mg/kg BW diluted in sterile saline. Control substance was IgG at the same concentration.

Axitinib (SigmaAldrich) was dissolved in DMSO and diluted with EtOH to 30 mg/mL. The solution was further diluted 8-fold with 10 % hydroxypropyl-β-cyclodextrin in saline to 3.75 mg/mL and applied at 25 mg/kg BW by i.p. injection. Control substance was DMSO/EtOH/hydroxypropyl-β-cyclodextrin in water following the same composition used for dissolving axitinib.

Combretastatin-A4 phosphate (fosbretabulin, CA4P,) was dissolved in PBS, sterile filtered and stored at -20 °C. In appropriate amount of the stock solution was further diluted prior to injection and applied in 100 µL saline at 25 mg/kg BW by i.p. injection. Control substance was 100 µL sterile saline (0.9% NaCl in water).

Tumor growth was followed by measuring perpendicular diameters of the tumors with a vernier caliper. Tumor volume was calculated using the equation V = π/6 x l x w^2^. In addition, tumors were excised post mortem and weighted. Only tumors that could be excised completely without additional invaded tissue were used for weight measurements.

### IHC and IF staining of tumor sections

H&E, IHC and IF staining was performed using standard techniques on formalin fixed paraffin embedded sections. Tissues for quantitative evaluation were processed in parallel. For quantification whole tissue sections were imaged on a Keyence BD 6000 microscope with an automated stage using a Nikon 10x objective. The individual images were stitched using the Keyence Analyzer software, to obtain a virtual slide. The whole virtual slide was used for quantification using the ImageJ software package (*rsbweb.nih.gov/ij/*).

Antibodies used for IHC, or IF: Cleaved caspase-3 (Cell Signaling Technology Cat# 9661, RRID:AB_2341188), carbonic anhydrase IX (Santa Cruz Biotechnology Cat# sc-25599, RRID:AB_2066539)), CD31 (Santa Cruz Biotechnology Cat# sc-28188, RRID:AB_2267979), CD34 (Abcam Cat# ab8158, RRID:AB_306316), Ki67 (Abcam Cat# ab16667 RRID:AB_302459) and anti-collagen IV (Bio-Rad, Rabbit, Cat# 2150-1470, RRID:AB_2082660).

### Production of lentivirial particles and generation of 4T1 and LLC cell lines expressing firefly luciferase

Lentiviral particles were generated in HEK 293T cells by co-transfection of the lentiviral vector pLVX-luc-IRES-puro (Clontech, Mountain View, CA) with the pCMV-dR8.9 and pCMV-VSV-G plasmids (both obtained from Addgene, Cambridge, MA), using a standard CaCl2-based transfection method(14). Supernatant collected from the HEK 293T cells 48h past transfection, was passed through a sterile 0.45 µm syringe filter and used to transfect 4T1 and LLC tumor cells after addition of sterile polybrene-solution (Hexadimethrinbromid, Sigma-Aldrich, 8 µg/mL final conc. In supernatant). This procedure was repeated once, 24 h later. Stable cells (4T1-fluc and LLC-fluc) were selected by treatment with puromycin (5 µg/mL) for 48h.

### 3D-angiography

For 3D-angiography, 50 µL of fluorescein-labeled dextran stock solution (Sigma, 20 mg/mL in 0.9% NaCl) and 50 µL of Dylight 648-labeled Isolectin GS-B4 (Nordic-MUbio, Susteren, The Netherlands. 500 µg/mL in 0.9% NaCl) were injected i.v. into tumor bearing mice. At the same time the mice were treated with PTX (20 mg/kg BW) or vehicle. Mice were sacrificed at different time points. 20 min before sacrificing the animals, they were injected with 50 µL of Dylight 594-labeled Isolectin GS-B4 (500 µg/mL in 0.9% NaCl). Tumors were removed and immersion fixed in 4 % PFA for 24 h. The tumors were cut in small cubes of approximately 5 mm, and subjected to dehydration by immersion in buffers with a gradually ascending ethanol concentration (50 %, 70%, 90% and 96% EtOH in TrisHCl, 50 mM, pH9.0). Remaining water was removed over 48 h in two changes of EtOHabs. before the samples were cleared over an addition 48 h in two changes of ethyl cinnamate (15). Images were acquired at high resolution (voxel size 86.9 nm x 86.9 nm x 713 nm) on a Leica SP5 LSCM.

#### Image processing and analysis

Image quality was first improved by deconvolution using the Huygens software suite (Scientific Volume Imaging, Hilversum, NL, https://svi.nl/). For deconvolution the given objective parameters (HCX PL APO CS 20.0x 0.70 DRY UV), resolution (0.0869 x 0.0869 x 0.713 µm) and refraction index of ethyl cinnamate (1.558) were used. Next images were processed using the ImageJ “Median 3D” filter and bicubic downsampled in x-y directions by the factor 2 to improve downstream image handling. Detailed analysis was done in Imaris 9.2.1 (Bitplane, Zurich, CH). Using constant threshold and filtering parameters surface rendering of the blood vessels in both channels (DyLight594 and DyLight 647) were generated and their volume calculated. The amount of FITC-dextran and Hoechst 33342 in the tissue was determined accordingly.

### CTC quantification

C57/Bl6J mice were implanted with LLC cells that stably expressed firefly luciferase and a selection marker for puromycin resistance (LLC-fluc). When tumors reached a size of 250 mm3 the mice were treated with PTX (20 mg/kg BW), mG6-31 (6 mg/kg BW) or a combination of both. Animals were sacrificed by CO2 asphyxiation 48 h later, and blood (> 200 mg) was immediately drawn from the right ventricle using heparinized syringes. Cells were collected by centrifugation, washed once with PBS and once with red blood cell (RBC) lysis buffer. After two more washes with PBS to remove the RBC lysis buffer, the cells from each sample were re-suspended in 3 mL DMEM with 10% FBS and Pen/Strep and plated into 30 wells of a 96-well MWD. The cells were cultivated for seven days under selective conditions (5 µg/mL puromycin for 48 h), lysed with 50 µL of passive lysis buffer (PLB, Promega, Mannheim, Germany) and stored at -80 °C. Luciferase activity was measured using the *luciferase assay system* from Promega, by combining 30 µL of cell lysate with 50 µL of luciferin solution. The resulting signal was recorded on a Tecan plate reader. Signals at least higher than average by at least two-times the standard deviation were counted as positive. CTCs per blood weight were calculated from the number of positive reads versus the amount of blood originally drawn.

### EC monolayer disruption under sheer stress

2.5 x 10^5^ HUVEC (< 6th passage) in 120 µL EGM-2 media (Lonza) were seeded in channel slides (µ-Slides 0.4 Luer, Ibidi, Martinsried, Germany) corresponding to a seeding density of 105 cells/mm2 and incubated at 37 °C. After 4 h an additional 90 µL EGM-2 were added to the reservoirs. Cells were left to attach overnight. The necessary number of individual chamber slides for the experiment (up to four) was connected in row with sterile silicone tubing via luer adapters. The assembled slides were then connected to the pump system (Ibidi, Martinsried, Germany), the whole fluidic system filled with 15 mL EGM-2 and flow adjusted to 2.4 mL/min. After 24 h under flow the flow rate was increased to 7.6 mL/min corresponding to a sheer stress of app. 7.6 dyn/cm. HUVEC were allowed to adjust to the sheer stress for 48 h with regular examination of the flow rate. After the adjustment phase reagents (PTX or CA4P) were added in 1 mL EGM-2 at an appropriate dilution to the pump reservoirs to reach the desired final concentration (50 nM). During the whole treatment period flow was kept constant at 7.6 mL/min. Channel slides and the pump system were kept in an incubator at all times to ensure constant temperature. Slides were removed from the flow system at the indicated time points and cells immediately fixed with ice-cold methanol, stained for VE-cadherin and F-actin using an Alexa-488-labeled phalloidin (Thermofisher) and counterstained with DAPI. Per timepoint four high-resolution images were acquired (Leica SP8 laser confocal microscope, 40x, NA1.8 objective) and evaluated using ImageJ.

### Scanning electron microscopy

Tumor samples were cut in pieces of around 5mm length immediately after resection, briefly washed in cold PBS and fixed first in McDowell-Trump fixative (2% PFA, 2.5% GA in 0.1M phosphate buffer, pH 7.2), at 4°°C 24 hours, and then in Fix in 3% Glutaraldehyde (in 0.1M phosphate buffer, pH 7.2) at 4°C for an additional 24 hours. Samples were washed 3 times for 30 min in 0.1M phosphate buffer, pH 7.2 and postfixed for 4h in 1% OsO4 (in 0.1M phosphate buffer, pH 7.2) at r.t.. The tissue was washed (2x water dd) and dehydrated in ascending ethanol (35%, 50%, 75%, 95% EtOH, 2h ea). To remove all traces of water the samples were submerged in five changes of 100% EtOH (4x 8h + ON). Dehydrated samples were immersed in hexamethyldisilazane (HMDS)(2 x 60 min.), dried in vacuo and sputtered with Au/Pd (10nm). Images were acquired by scanning on a Tecan Mira3 microscope in resolution mode.

### Electric cell-substrate impedance sensing

For electric cell-substrate impedance sensing (ECIS) a protocol based on the method published by Szulcek *et al.* was used (16). In short, 8-well ECIS slides (8W10E+, Ibidi, Martinsried, Germany) were washed for 10 min with sterile 10 mM cystein in water and twice with sterile pure water. Afterwards early passage HUVEC were seeded in 500 µL EGM-2 at 4×10^5^ cells/well. The slides were incubated for 48 h at 37 °C, 5% CO2 until HUVEC were confluent. Cells were supplied with 580 µL of fresh EGM-2 and equilibrated for 5 h in the ECIS 16 well station (Ibidi) while measuring impedance in multi-frequency (MFT) mode. After 5 h measurement was paused and drugs added in 20 µL of EGM-2 before measurement was continued for > 24 h. Calculation of resistance was performed using the ECIS software.

### Tube formation assay

Tube formation assays were performed on angiogenesis µ-slides (Ibidi, Martinsried, Germany). In short wells were coated with 10 µL matrigel (Corning Life Sciences, Amsterdam, The Netherlands). The matrigel was incubated for 30 min at 37 °C to solidify. HUVEC (< 6^th^ passage) were seeded with 4000 cells/well in 50 µL EGM-2. The Slides were again incubated for 18 h to let tube like structures form. The cells were treated with EGTA or 2-APB at concentrations of 1 mM and 75 µM respectively, before PTX was added at a final concentration of 50 nM. Cells were stained 10 min before imaging with Calcium AM, and images were acquired at 5x and 10x magnification using the standard fluorescein filter channel on a Leica fluorescence microscope. Images were analyzed without further manipulation using the Angiogenesis Analyzer tool for ImageJ (17, 18).

### RNA-isolation

RNA was isolated from cells using the RNeasy Kit (Qiagen, Hilden, Germany) according to the manufacturer’s recommendations.

RNA was isolated from fresh tumor samples using the Trizol reagent (Life Technologies, Darmstadt, Germany) according to the manufacturer’s recommendation.

### mRNA-Quantification

mRNA-expression levels were quantified using the GeXP-System (BeckmanCoulter, Krefeld, Germany). Protocols for reverse transcription, amplification, labeling, gel electrophoresis and quantification were used as recommended by the manufacturer. RNA-levels were normalized to levels of housekeeping genes β-2-microglobulin (B2M) and ribosomal protein S29 (RPS29)(19). Analysis was done with three technical replicates per biological sample. Mean values of technical replicates were used for statistical analysis.

### Patient data evaluation and gene set enrichment analysis (GSEA)

For details regarding study design, treatment schedule, regiment, sample preparation and microarray data processing see https://clinicaltrials.gov/ct2/show/NCT00773695 and (20). The data set is available at: https://www.ebi.ac.uk/arrayexpress/experiments/E-MTAB-4439/.

Matched patient expression profiles pre- and posttreatment were identified. This resulted in 58 and 53 matched profiles, respectively (in the CTX-alone and in the BEV+CTX groups). Expression changes that occurred in these matched profiles, presumably in response to therapy, were calculated and used to analyze differences in the response to the two different treatments.

GSEA was performed using the software available at https://www.gsea-msigdb.org/gsea/index.jsp using a signal-to-noise metrics and a weighted enrichment statistic (21, 22). In addition to self-curated gene sets for EMT, invasiveness, angiogenesis and hypoxia (Supplemental Table 1), GeneOntology gene sets of biological processes (GOBP) and the Hallmark gene sets maintained and available at https://www.gsea-msigdb.org/gsea/msigdb/genesets.jsp?collection=H were used to test for differences between the treatment groups.

#### Association with survival data

The NCT00773695 dataset had transcriptomic profiles of 132 BRCA patients at pre-treatment, and post-treatment. We made three comparisons of BRCA patient transcriptomic profiles from this data; (I) bevacizumab + chemo post-treatment vs pre-treatment, (II) only chemo post-treatment vs pre-treatment, (III) bevacizumab + chemo post-treatment vs only chemo post-treatment. From these comparisons treatment-specific genes whose expressions were upregulated or downregulated in each treatment arm compared to the other arms or pre-treatment were identified. The gene signature upregulated and downregulated in bevacizumab + chemo post-treatment vs pre-treatment comprised of 60 genes and 34 genes respectively. The gene signature upregulated and downregulated in only chemo post-treatment vs pre-treatment comprised of 90 genes and 126 genes respectively. The gene signature upregulated and downregulated in bevacizumab + chemo post-treatment vs only chemo post-treatment comprised of 218 genes and 104 genes respectively. We performed single-sample geneset enrichment analysis (ssGSEA) on the upregulated and downregulated genesets from the above three comparisons. To correlate the single-sample enrichment of these genesets with breast cancer patient survival from TCGA, we extrapolated the treatment-specific upregulated/downregulated genesets from the above analysis into TCGA transcriptomic data and associated the ssGSEA enrichment scores of these genesets (calculated on TCGA BRCA transcriptomic data) with disease-free survival data of stage I and stage II BRCA patients from TCGA. For checking the association of the geneset enrichments with disease-free survival, BRCA patients are stratified into two groups, high enrichment of the gene signature (greater than 75th quantile in population) and low enrichment of the gene signature (less than or equal to 75th quantile in population). Kaplan-Meier analysis was performed for comparison of disease-free survival between these two stratified groups.

### Statistical Analysis

All statistical analysis was done using the Prism5 Software (GraphPad, LaJolla, CA). Differences between two groups were analyzed using an unpaired, two-tailed Student’s T-test. In parallel the samples were tested for significant variation of variance, and if necessary a Welch correction was included in the statistical analysis. For statistical analysis of metastatic incidence and size of metastases between groups the Mann-Whitney test was used, as a Gaussian distribution could not be assumed. All statistical tests were performed between sets of individual biological replicates.

## Results

We tested the effect of combining taxane-based chemotherapy with anti-angiogenic therapy in a mouse model of stage IV metastatic breast cancer: fully established tumors arising from orthotopically implanted 4T1 cells in syngeneic Balb/c mice were continuously treated with paclitaxel (PTX) and mG6-31, a monoclonal antibody active against murine VEGF-A (23). PTX by itself (administered at 20 mg/kg BW twice in a six-day cycle for four cycles)(12) had no significant effect on primary tumor growth (Fig. 1A, B). However, in combination with mG6-31 (at 6 mg/kg BW q6d)(24) PTX completely blocked growth of the aggressive tumors. The effect on lung metastasis was in opposition to the observation on the primary tumors: PTX alone significantly reduced the number of metastases, while mG6-31 alone, which was effective in reducing primary tumor growth, had no effect on lung metastases (Fig. 1C, D). Interestingly, adding mG6-31 to PTX treatment abrogated the beneficial effect of PTX on the rate of metastasis. Although, the size of lung metastases was significantly smaller in both the PTX and PTX + mG6-31 groups, the number of metastases formed was as high after PTX + mG6-31 combination treatment as in the control group (Fig. 1D, E).

**Figure 1:**
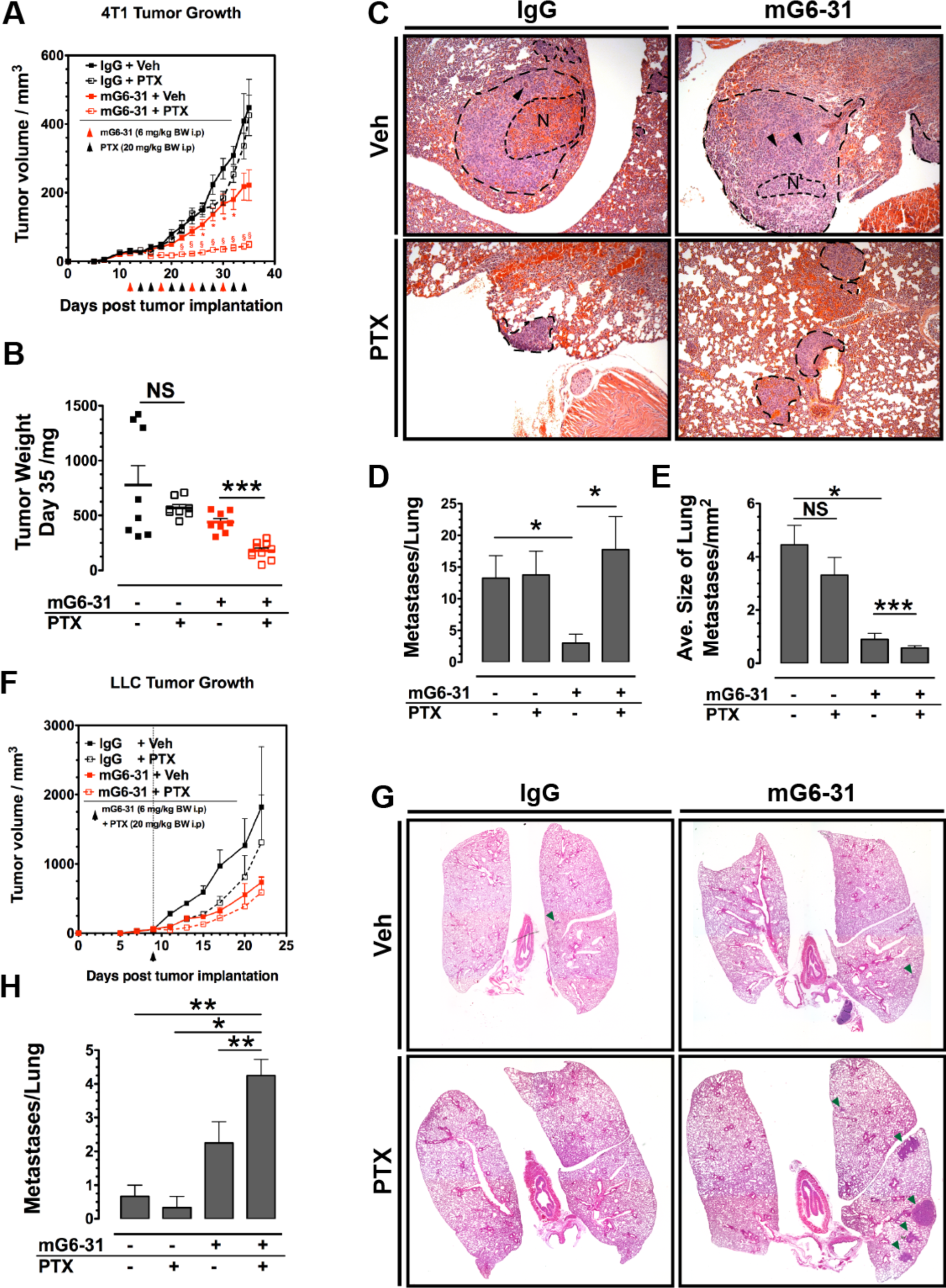
Combination of paclitaxel and anti-VEGF treatment increases rate of metastasis. (A) Growth of primary 4T1 breast adenocarcinomas treated with PTX (20 mg/kg BW, i.p.), mG6-31 (6 mg/kg BW, i.p.) or a combination of both. The combination of PTX and mG6-31 suppressed growth of the tumors effectively, PTX alone had no significant impact on primary tumor growth. Days when treatment was applied are indicated (black and red arrow heads). (B) Weight of treated 4T1 primary tumors 35 days post implantation. (C) H&E staining of lungs from 4T1 tumor bearing mice after treatment. Metastases are marked. Scale bar = 100 µm. (D) Numbers of metastases found in lungs from 4T1 tumor bearing mice after treatment. (E) Size of metastases found in lungs from 4T1 tumor bearing mice after treatment. (F) Growth of primary LLC tumors treated with a single dose of PTX (20 mg/kg BW, i.p.), mG6-31 (6 mg/kg BW, i.p.) or a combination of both. Drugs were administered on day 9 post tumor implantation, when tumors reached a size of 50 mm^3^. (G) Whole mount images of sections from H&E-stained lungs from LLC tumor bearing mice after treatment. Metastases are indicated (green arrow heads). (H) Numbers of metastases found in lungs from LLC tumor bearing mice after treatment. Error bars: ± SEM. Asterisks indicate statistical significance versus control, *: P < 0.05, **: P < 0.01, ***: P < 0.001.

PTX is very effective against tumor cells in the circulation (CTCs) and nascent tumors at distal sides, as these cells are not protected by an established tumor microenvironment (TME)(25). Thus, we hypothesized that the metastasis promoting effect of combining PTX with mG6-31 is partially concealed by the continuation of PTX treatment, which kills disseminating CTCs that would otherwise form additional metastases. Therefore, we treated tumor-bearing mice with a single round of concomitantly administered PTX + mG6-31 and waited two weeks before assessing the impact of this single treatment on lung metastasis. This schedule mirrors how these drugs are applied in patients (3-5). For this experiment the lewis lung carcinoma (LLC) model was chosen as it has a lower basal metastasis rate as the 4T1 model. Treatment again affected primary tumor growth, although as the treatment was not repeated the effect did not persist (Fig. 1F). PTX + mG6-31 strongly increased the number of lung metastases compared to non-treated, PTX-alone or mG6-31-alone mice (Fig. 1G, H).

To evaluate if the observed effect could also be found with other anti-angiogenic drugs we repeated the single treatment study in the breast cancer model, however substituting axitinib (AXI), a small molecule TKI targeting the VEGF-receptor family, for mG6-31. Moreover, the short serum half-life of AXI allowed also to reverse the order of application in a three-day schedule: tumor-bearing animals were treated with one round of PTX followed by two injections of AXI (25 mg/kg BW i.p., PTX→AXI)(26) on the two following days or with two injections of AXI followed by one injection of PTX on the third day (AXI→PTX). Controls received vehicle, a single injection of PTX or three injections of AXI. Both combination treatments significantly reduced growth of primary tumors (Sup. Fig. S1A). However, lungs resected 11 days after initiation of treatment, showed a substantial increase of metastatic lesions only in the PTX→AXI group, while reversing the order (AXI→PTX) had no effect on lung metastasis (Sup. Fig. S1B, C).

To determine the immediate effect of taxane-based therapy in combination with anti-angiogenic agents on the TME, we analyzed primary tumors 48 h post therapy for changes in vascularization and for indications of reduced supply. 4T1 and LLC tumors were treated with a single injection of PTX, mG6-31 or a combination of both and resected 48 h later. In 4T1 tumors effects were not apparent in the single therapy arms within that short period after treatment. However, combination of both PTX and mG6-31 significantly reduced the number of CD31^+^ endothelial cells (ECs, Sup. Fig. S2A), and caused a strong increase of hypoxia, indicated by elevated expression of carbonic anhydrase IX (CAIX, Sup. Fig. S2B, C). In addition, the tumors that underwent combination treatment also showed increased central necrosis, further indicating either reduced supply or a synergistic increase of cytotoxicity (Fig. 2A, B). In LLC tumors PTX/mG6-31-treatment had similar consequences after 48h. However, in LLCs mG6-31 was able to reduce vascularization and cause hypoxia as stand-alone therapy (Sup. Fig. S2A-C). PTX alone had a less pronounced effect and values did not reach statistical significance. In combining both drugs, the anti-vascular and hypoxia-inducing effect was strongly enhanced. After PTX treatment the vast majority of the tumor already appeared necrotic (79.4 ± 8.7% of tumor area), an effect not further enhanced by addition of mG6-31 (Fig. 2A).

**Figure 2:**
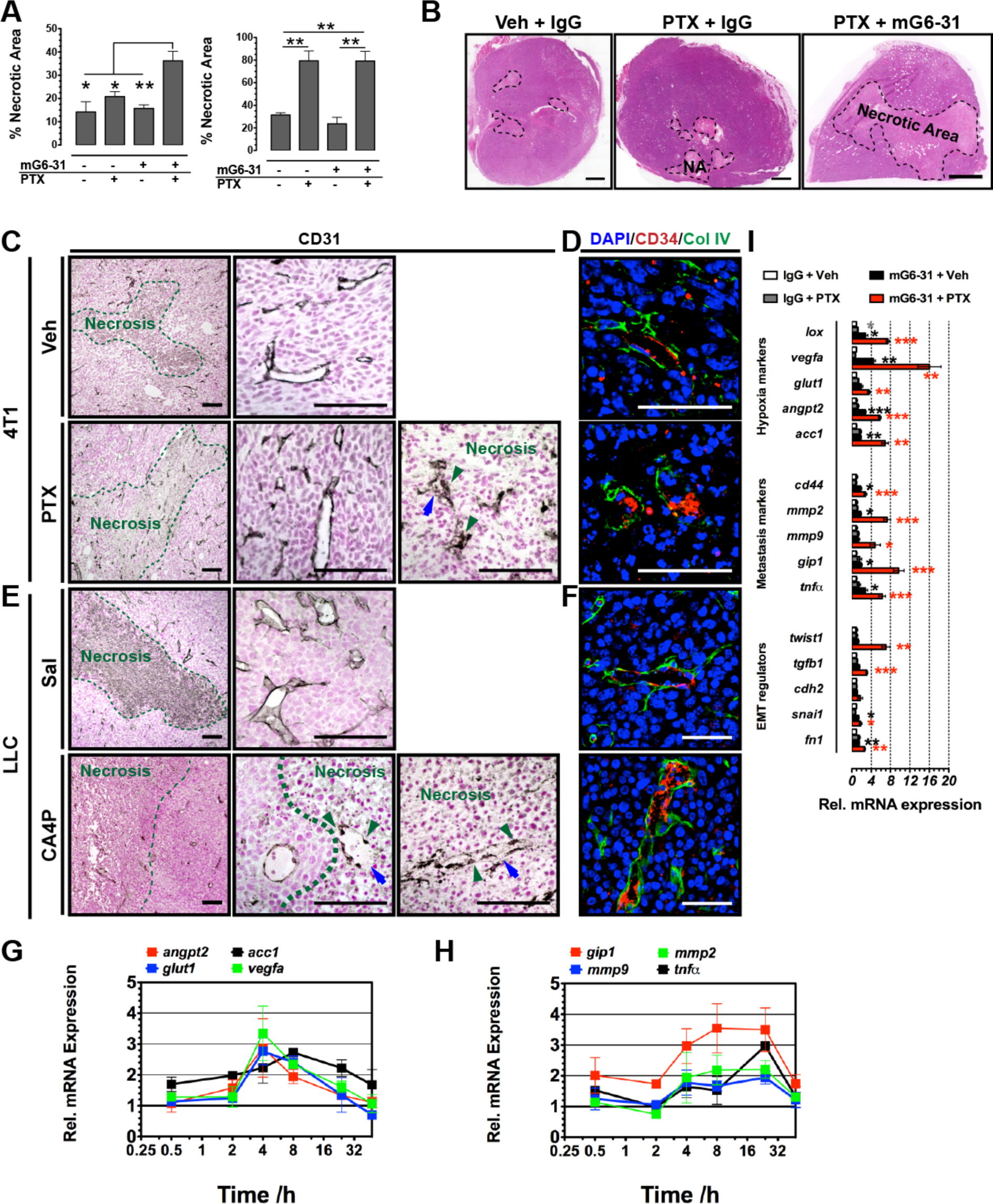
Combination of paclitaxel and anti-VEGF treatment leads to acute hypoxia. **(A)** Quantification of necrosis in 4T1 and LLC tumors 48 after treatment with PTX, mG6-31 or a combination of both. Whole H&E-stained tumor sections were evaluated. **(B)** Images of whole H&E-stained LLC tumor sections after treatment with PTX or PTX + mG631. Scale bar = 1 mm. **(C)** Images of 4T1 tumors 48 after treatment with PTX. After treatment CD31 staining shows blood vessel structures in the necrotic areas of the tumors, which is absent from untreated control tumors. The vessels structures within the necrosis are characterized by rounded endothelial cells (green arrow heads) and large portions of the vessel rim no longer covered by the endothelium (CD31^-^, blue arrows). Scale bar = 100 µm. **(D)** Confocal images of 4T1 tumors 48 after treatment with PTX, stained for CD34 and the basal lamina component collagen IV. After treatment CD34^+^ endothelial cells in the necrotic areas appear detached from the basal lamina. Scale bar = 100 µm. **(E)** Images of LLC tumors 48 after treatment with CA4P (25 mg/kg BW). After treatment CD31 staining shows blood vessel structures in the necrotic areas of the tumors, which is absent from untreated control tumors. The vessels structures within the necrosis are characterized by rounded endothelial cells (green arrow heads) and large portions of the vessel rim no longer covered by the endothelium (CD31^-^, blue arrows). Scale bar = 100 µm. **(F)** Confocal images of LLC tumors 48 after treatment with CA4P (25 mg/kg BW), stained for CD34 and the basal lamina component collagen IV. After treatment CD34^+^ endothelial cells in the necrotic areas appear detached from the basal lamina. Scale bar = 100 µm. **(G)** Time course of relative mRNA expression levels of markers for hypoxia and metabolic stress in LLC tumors within 48 h past PTX (20 mg/kg BW) treatment (n=4). **(H)** Time course of relative mRNA expression levels of markers for invasiveness stress in LLC tumors within 48 h past PTX (20 mg/kg BW) treatment (n=4). (I) Relative mRNA expression levels of markers for hypoxia and metabolic stress, metastatic behavior or EMT in 4T1 tumors 48 after treatment with PTX, mG6-31 or a combination of both (n=4). Asterisks indicate statistical significance versus control, *: P < 0.05, **: P < 0.01, ***: P < 0.001.

To better understand the effect that taxanes exert on tumor vessels, we histologically examined the vasculature of tumors 48 h post treatment with PTX in more detail. There was a significant difference observed in the necrotic regions of treated tumors compared to non-treated controls: In 4T1 control tumors the necrotic cores of the tumors were devoid of any remnants of blood vessels (Fig. 2C, D). In contrast, 48h after PTX treatment the necrotic fraction of the tumors was strongly increased and in these recently damaged areas blood vessels are clearly discernable with a microvascular density (MVD) similar to the surrounding non-necrotic parts. However, the vessels display strongly rounded endothelial nuclei and disintegrated luminal endothelial lining. In stark contrast the endothelial layer in perinecrotic areas appeared normal, not affected by the treatment. In LLC tumors the effect of PTX was analogous (Sup. Fig. S3A, B).

These results indicated that PTX and other taxanes have not an angiogenesis inhibiting effect but act like vascular disrupting agents (VDAs), causing a rounding of endothelial cells (ECs), disruption of vascular integrity, and eventually vascular collapse leading to reduced supply and tumor necrosis. We therefore, tried to reproduce the results substituting the VDA combretastatin-A4 phosphate (fosbretabulin, CA4P, applied at 25 mg/kg BW) for PTX. Indeed, the combination treatment with CA4P and mG6-31 caused a significant increase in metastasis (Sup. Fig. S4A-C). Moreover, 48 h after treatment with CA4P the tumors displayed identical vascular damages as observed at this time point after application of PTX (Fig. 2E, F).

The rapid collapse of the tumor vasculature leading to undersupply of the surrounding tumor tissue caused by PTX, should be reflected in expression changes of response genes. Indeed, mRNA-expression of various genes responsive to hypoxic (*angpt2, glut1* and *vegfa*,) and metabolic stress (*acc1*) significantly increased in LLC tumors within 4 hours after PTX application (Fig. 2G). Markers of increased metastatic and invasive behavior (*gip1*, *mmp2*, *mmp9* and *tnfα*,) also increased but their levels peaked later, 8h to 24h post treatment (Fig. 2H). 48h post treatment with PTX expression levels of markers of undersupply and metastasis were widely returned to normal in both 4T1 and LLC tumors. However, when animals were treated with mG6-31 concurrently to PTX mRNA levels of hypoxia, metastasis and EMT markers remained high, indicating protracted hypoxic stress (Fig. 2I).

To test whether combining taxanes with antiangiogenic therapy has a similar effect on the metastatic and hypoxic profile in patients, we evaluated microarray data sets from the NCT00773695 trial, that followed the mRNA-expression levels in Her2-negativ breast cancer patients undergoing neoadjuvant taxane-based therapy (CTX) with or without bevacizumab (20). Addition of BEV resulted in significant expression changes in a much wider range of genes than CTX alone (Sup. Fig. S5A, B). We first used gene set enrichment analysis (GSEA) to compared the differences between both treatment groups with respect to the expression changes from pre-treatment biopsies and biopsies taken after the start of the taxane therapy. A wide range of externally curated gene sets - among them the gene ontology sets for biological processes (GOBP) - and self-compiled sets for positive regulators and indicators of invasiveness, hypoxia, angiogenesis and EMT were tested (Sup. table 1). Results demonstrated that expression of genes indicative of invasiveness, hypoxia, angiogenesis and potentially metastatic behavior increased much stronger when patients were treated with the CTX+BEV combination than when they were treated with CTX alone (Fig. 3A-F, Sup. Fig. S5C, D)). We next evaluated expression changes *within* the two patient groups between pre-treatment biopsies and biopsies taken after start of the taxane treatment. In the CTX alone group only angiogenic signaling was significantly enhanced (Sup. Fig. S5E), while treatment with a combination of CTX+BEV increased also hypoxia and invasive behavior (Sup. Fig. 5F). Thus, the effects of a combined taxane plus anti-angiogenic therapy in patients reflects our observation in mice. Moreover, taxane-based therapy alone increases angiogenic signaling in breast cancer patients, an addition of BEV leads then to hypoxia and invasiveness (Sup. Fig. S5G). Next, we examined whether the transcriptional changes observed after treatment were also correlated to outcome. Based on the NCT00773695 trial data we made three comparisons of BRCA patient transcriptomic profiles from this data; (I) CTX+BEV post-treatment vs. pre-treatment, (II) CTX only post-treatment vs pre-treatment, (III) CTX+BEV post-treatment vs CTX only post-treatment. From these comparisons treatment-specific genes with altered (up or down) expression between arms were identified (Fig. 2G). Single-sample GSEA (ssGSEA) on the altered genesets was performed and extrapolated into TCGA BRCA transcriptomic data. Finally, the ssGSEA enrichment scores calculated on TCGA BRCA transcriptomic data was stratified into two groups, high and low enrichment of the gene signature (> 75^th^ and ≤ 75^th^ quantile in population, respectively), and associated with disease-free survival (DSF) data of stage I + II TCGA BRCA patients. Kaplan-Meier analysis demonstrated that high enrichment of the gene signature upregulated in CTX+BEV post- vs pre-treatment tends to associate weakly with better DFS (p = 0.16, Fig. 2H), while low enrichment of the gene signature downregulated in CTX+BEV post- vs pre-treatment (p-value = 0.04, Fig. 2I). On the other hand, enrichment of the gene signature up- or downregulated in CTX+BEV vs CTX only did not associate with DFS (p = 0.49 and 0.69, respectively, Fig. 2J,K). Thus, the analysis indicated towards a treatment benefit of CTX+BEV in terms of DFS but no apparent improvement could be inferred compared to CTX alone.

**Figure 3:**
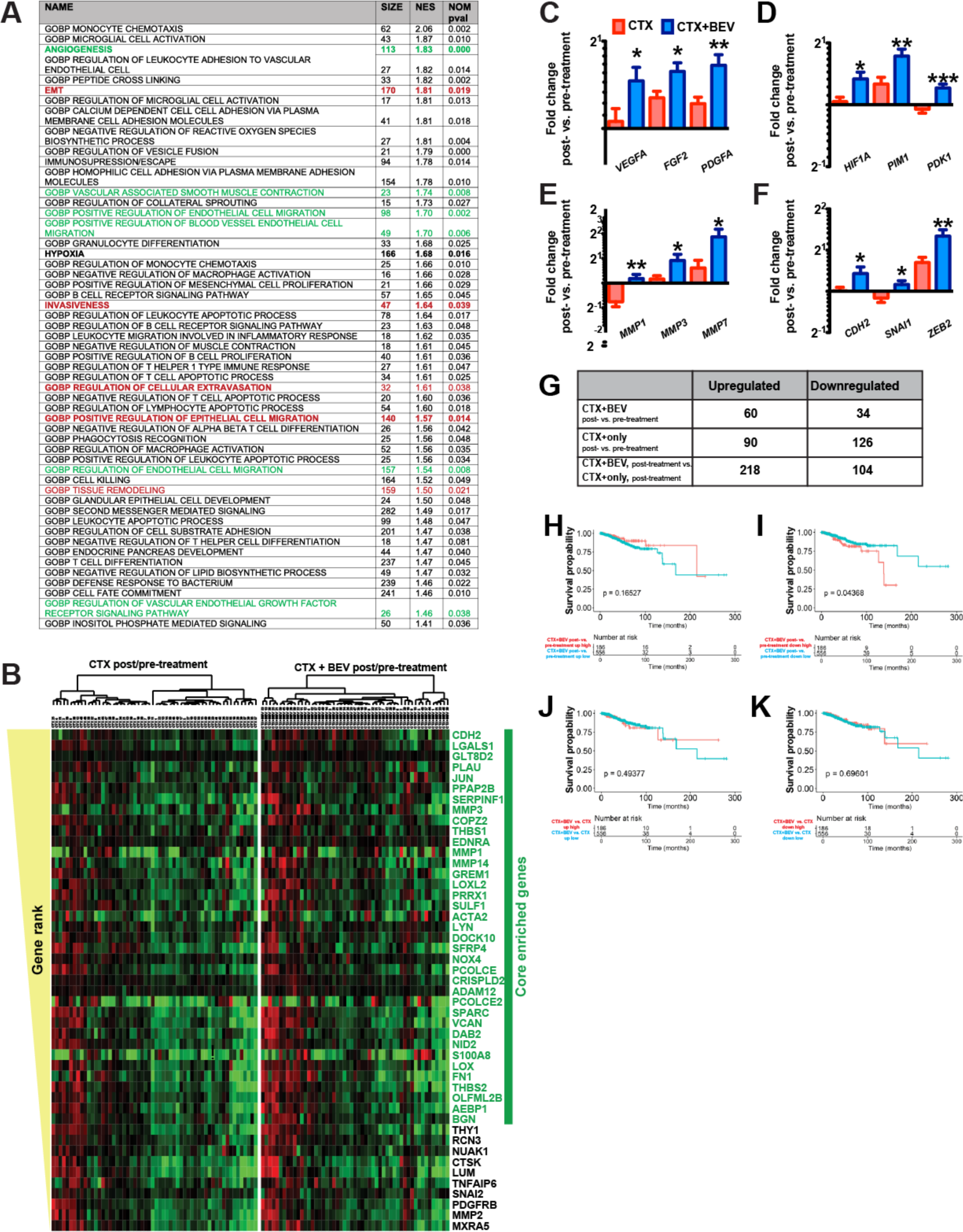
Combination of taxanes with bevacizumab increases expression of genes involved in hypoxia response, invasiveness and EMT in human patients. (A) Selection of gene sets that were significantly (P < 0.05) stronger changed in the CTX+BEV group vs the CTX group between the pre- and posttreatment biopsies. Self-curated gene sets are accentuated in bold face, gene sets are shown in red, sets angiogenesis in green. GOBP: gene ontology sets with respect to defined biological processes. (B) Normalized expression changes in genes of the invasiveness gene set in both treatment groups post- vs. pretreatment. Genes are sorted according to rank. **(C)** Expression changes of selected genes involved in angiogenesis in both treatment groups post- vs. pretreatment. **(D)** Expression changes of selected genes involved in hypoxic response in both treatment groups between post- vs. pretreatment. **(E)** Expression changes of selected genes encoding metalloproteases involved in cell invasion in both treatment groups between post- vs. pretreatment. **(F)** Expression changes of selected genes involved in epithelial-to-mesenchymal transition in both treatment groups post- vs. pretreatment. **(G)** Extrapolating the treatment-specific gene signatures from clinical trial to TCGA cohort and associating with disease-free survival for stage I and stage II BRCA patients. High enrichment of the gene signature upregulated in CTX+BEV post- vs pre-treatment (p-value = 0.16). **(H)** Low enrichment of the gene signature downregulated in CTX+BEV post- vs pre-treatment (p-value = 0.04). **(I)** Enrichment of the gene signature upregulated in CTX+BEV vs CTX only. **(J)** Enrichment of the gene signature upregulated in CTX+BEV vs CTX only. Neither up- nor downregulated GS associate with DFS. **(K)** Asterisks indicate statistical significance versus control, *: P < 0.05, **: P < 0.01, ***: P < 0.001. Asterisks indicate statistical significance versus control, *: P < 0.05, **: P < 0.01, ***: P < 0.001.

The effect of taxanes on endothelial cells and the vasculature is not sufficiently examined. To get time-resolved information about the effects of PTX on endothelial barrier function electric cell-substrate impedance sensing (ECIS) was used. HUVEC were grown to confluency in ECIS chambers and subjected to treatment with PTX or CA4P at 50 nM or 250 nM. Analysis of the primary data obtained by multi-frequency impedance scanning yielded resistance as a measure of barrier function (Rb) and a quantitative description of cell shape (α) (16). PTX exposure resulted in a rapid reduction of Rb indicating a breakdown of the endothelial barrier (Fig. 4A). Interestingly, the effect was stronger at the lower concentration of 50 nM than at 250 nM. At 50 nM Rb reached a low after 1.7 h (36 % of the normalized Rb). CA4P showed an overall stronger and dose correlated effect. Two major differences could be observed between PTX and CA4P effects on endothelial monolayers: the PTX-triggered breakdown partially regressed, although the drug was not removed from the media. Furthermore, the strong increase in α caused by PTX indicated a higher resistance through the ECs, potentially caused by a rounding of the ECs (Fig. 4B). In contrast, CA4P treatment resulted after a brief recovery phase in a decline towards total loss of the barrier function, and a concomitant decrease in α. Docetaxel (DTX), another taxane, showed the same effect as PTX and, again a concentration of 50 nM was more effective in breaking the EC barrier than 250 nM (Sup. Fig. S6A). A wide range of other commonly used but non-tubulin binding chemotherapeutics had no significant effects on the EC layer (Sup. Fig. S6B). Monolayers of fibroblasts (NHDF) did not display significant effects after treatment with either PTX or CA4P (Fig. S6C). To more closely mimic the situation *in vivo*, we cultivated HUVEC in channel slides to confluency and subjected them to increasing shear stress by applying a flow of culture media. After cells were adjusted to a shear stress of 7.5 Dyn/cm^2^ for 24 h PTX or CA4P was added at a concentration of 50 nM. 30 min after PTX-treatment a dense pattern of large holes appeared in the EC-monolayer, caused by an apparent uniform contraction of the individual ECs (Fig. 4C, E, F). 8h after PTX-treatment started these openings in the monolayer had closed again and stable EC-EC junctions had been reestablished as shown by VE-cadherin staining. The effect of CA4P on the other hand occurred significantly later and was not reversible (Fig. 4D, E, F). While ECs treated with PTX underwent a simple morphological modification, CA4P-treated ECs appeared to be terminally damaged by severe toxic stress.

**Figure 4:**
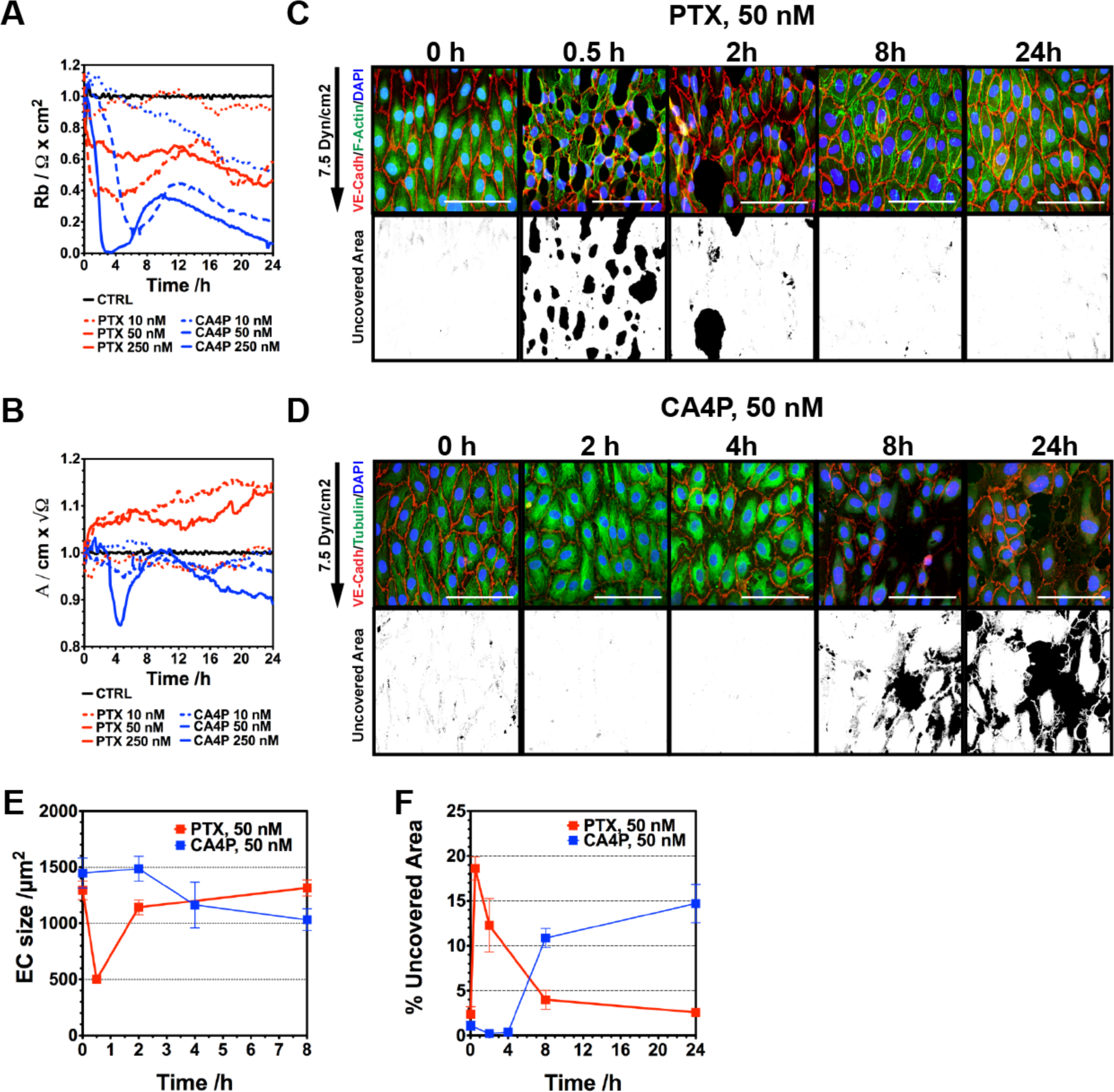
Paclitaxel treatment at nano-molar doses leads to rapid, reversible breakdown of the endothelial barrier. **(A)** Resistance (Rb) of the endothelial barrier after treatment with PTX or CA4P at 50 nM and 250 nM. Results of ECIS measurement on HUVEC. Reduction of Rb indicates impaired endothelial barrier, that is after PTX-treatment reversible. **(B)** Alpha (α) as a measurement of trans-endothelial resistance. Results of ECIS measurement on HUVEC. PTX and CA4P have different effects on cell shape resulting in different changes of α. **(C)** Time course images of endothelial monolayers treated with 50 nM PTX under flow. In the lower row openings in the endothelial monolayers are displayed as black areas not covered by cells. Scale bar = 10 µm. **(D)** Time course images of endothelial monolayers treated with 50 nM CA4P under flow. Changes in tubulin distribution are already present after 2h. However, only after 8h tubulin has been dissolved and the effect on endothelial barrier is apparent. In the lower row openings in the endothelial monolayers are displayed as black areas not covered by cells. Scale bar = 10 µm. **(E)**Time course evaluation of endothelial shape in monolayers treated with 50 nM PTX or 50 nM CA4P under flow. **(F)**Time course evaluation of openings in endothelial monolayers treated with 50 nM PTX or 50 nM CA4P under flow.

As the permeabilization of the EC monolayer by PTX occurs rapidly it is unlikely to be an effect exerted through the canonical tubulin-stabilization mechanism. A faster mechanism has to trigger the contraction of the ECs. Moreover, the ECs in the confluent monolayer are not proliferating and therefore widely resistant to the cytotoxic effects of PTX, again indicating an alternative mechanism (Sup. Fig. S7A-E).

Although, the EC-contraction in response to PTX is reversible *in vitro*, *in vivo* the resulting permeabilization of the endothelium should lead to serum efflux and a vascular collapse that is largely irreversible. We treated tumor-bearing mice (4T1 or LLC) with PTX and removed tumors at various time points for evaluation of the vasculature via SEM and 3D angiography. In SEM images of resected tumors fractures appeared on the surface of blood vessels 45 min after PTX application (Fig. 5A, B). Number and severity of the observed breaks increased over time.

**Figure 5:**
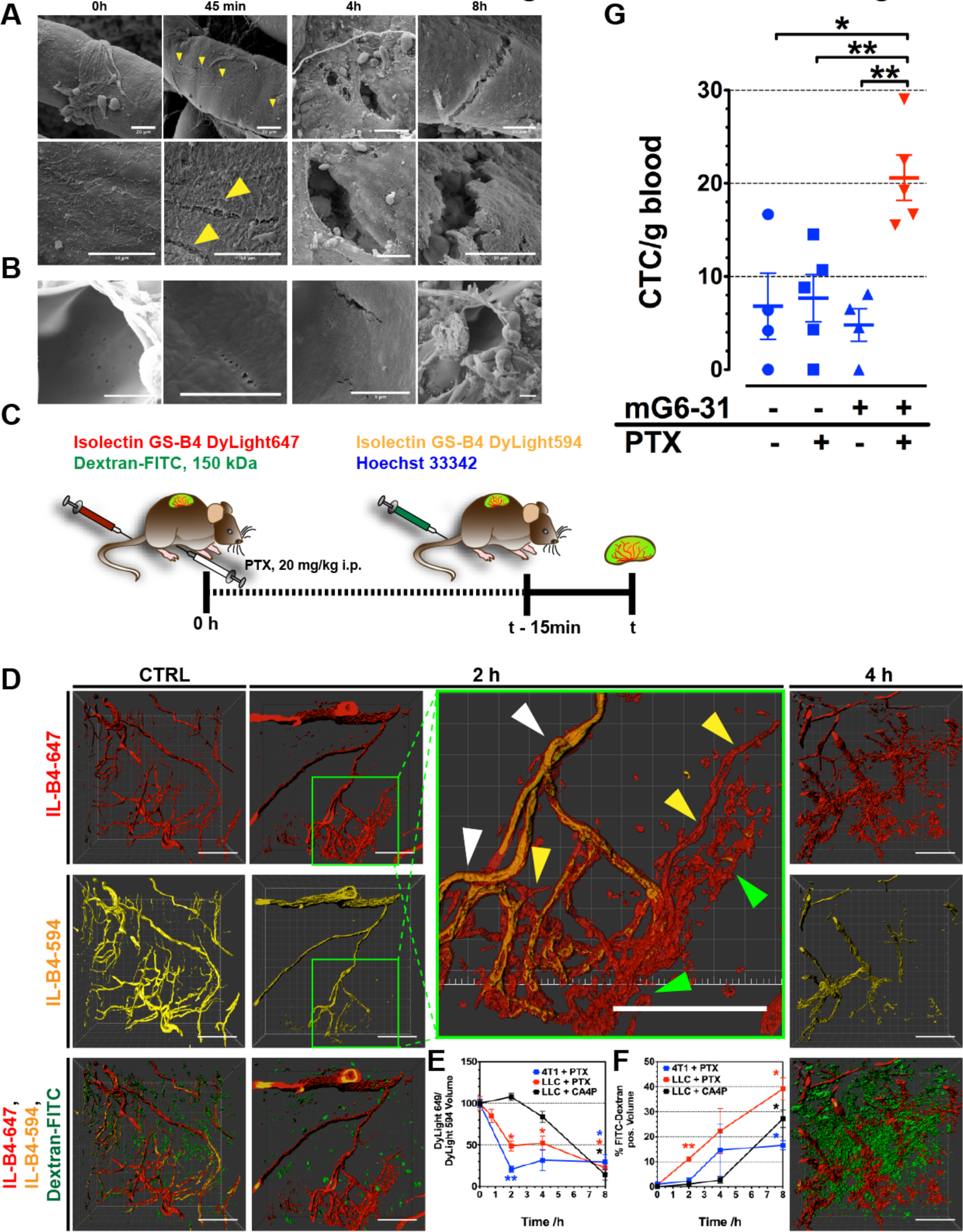
Paclitaxel treatment causes rapid vascular collapse. **(A)** SEM images of LLC tumor blood vessels at different time points after application of PTX (20 mg/kg BW). Scale bar = 20 µm (upper row) and 10 µm (lower row). **(B)** SEM images of 4T1 tumor blood vessels at different time points after application of PTX (20 mg/kg BW). Scale bar = 10 µm. **(C)** Schematic schedule for the application of tracer substances in the vessel collapse experiments. IL-B4-DyL647 was applied immediately before treatment with PTX (20 mg/kg BW) or CA4P (25 mg/kg BW), labeling the perfused vascular network at t0 in the unperturbed tumor. IL-B4-DyL594 was applied 15 min before necropsy (to allow for even distribution), labeling only blood vessels still intact. **(D)** 3D images of vascular networks in 4T1 tumors after PTX application. Differences in IL-B4-DyL647 and IL-B4-DyL594 signals show vascular collapse. After 2h some of the initially perfused vessels are still perfused (white arrow heads), others are not-longer supplied but show a still intact structure (yellow arrow heads), while a third class is completely disintegrated (green arrow heads). Scale bar = 100 µm. **(E)** Ratio between IL-B4-DyL647^+^ and IL-B4-DyL594^+^ vessel volume indicating the degree of vascular collapse between t0 and time of tissue collection. **(F)** Percentage of tissue volume infiltrated by HMW dextran demonstrating increased vascular permeability in response to PTX and CA4P. **(G)** Number of circulating tumor cells in LLC tumor bearing mice 48 h post treatment with PTX, mG6-31 or a combination of both. Asterisks indicate statistical significance versus control, *: P < 0.05, **: P < 0.01, ***: P < 0.001.

For 3D angiography two isolectins labeled with different fluorophores were applied to tumor-bearing animals subsequently: Dylight 649-labeled isolectin-B4 (IL-B4-649) was injected at the time of PTX-treatment and Dylight 594-labeled isolectin-B4 (IL-B4-594) later, 15 min before necropsy (Fig. 5C). Signal of the two dyes should indicate perfused vessels at the different time points, and thereby allow to measure the extend of changes in perfusion in response to therapy. In addition, we injected FITC-labeled HMW-dextran (500 kDa) together with PTX to visualize leakage. The analyzed 3D-data showed a rapid decrease in vessel perfusion that was already significantly reduced 45 min after PTX application (Fig. 5D, E; Sup. Fig. S8A). In contrast to the situation *in vitro*, a rebound was not observable, and perfusion decreased steadily over time as more and more vessels collapsed. In parallel, the amount of FITC-dextran increased (Fig. 5D, F; Sup. Fig. S8A). The tracer Hoechst 33342 was injected in LLC bearing mice together with IL-B4-594 15 min before necropsy, indicating the regions of the tumor accessible by small drug-like molecules (Fig. 5C). After PTX-treatment the supplied part of the tumor successively reduced, reaching a minimum after 4h (Sup. Fig. S8B). When treated with CA4P the vasculature of LLC tumors responded similar, however with the expected delay when compared to the response towards PTX (Fig. 5E, F, Sup. Fig. S8C).

Finally, we tested the effect of a PTX/mG6-31 combination treatment on the number of circulating tumor cells (CTCs). Blood was drawn from animals bearing LLC tumors stably expressing firefly luciferase (LLC-*fluc*), and the blood samples evaluated for the number of *fluc*-expressing cells. 48 h post treatment the concentration of CTCs tripled in animals receiving PTX + mG6-31 (20.6 ± 2.4 vs. 6.8 ± 3.5 CTCs/g blood, Figure 5G, Sup. Fig. S9A-D). Treatment with PTX or mG6-31 alone did not result in a significant increase of CTCs in the peripheral blood.

## Discussion

Our results clearly show that PTX acts as a vascular disrupting agent, targeting with some selectivity the activated and defective vasculature of the tumor. Although, the effect of PTX on endothelial cells is reversible *in vitro*, the disruption of tight and adherens junctions leads *in vivo* to a strongly increased para-endothelial flux, thus leakage of serum into the tumor parenchyma and an irreversible vascular collapse. The EC rounding effect of PTX that leads to breakdown of the vascular barrier involves Ca^2+^ influx via TRP channels. Pretreatment of patients with a TRP inhibitor could therefore be used to protect the tumor vasculature from this VDA effect, if it turns out that this would be beneficial. As we have shown previously the acute hypoxic stress resulting from the rapid collapse, leads to a strong increase of pro-angiogenic signaling that i.a. mobilizes EPCs from the bone marrow(12). Inhibition of VEGF (or SDF-1) signaling blocks this vasculogenic rescue, further increasing hypoxia (Sup. Fig. S10). While this concerted attack on the tumor’s blood supply has indeed a profound effect on the viability of established, vascularized tumors it also increases the risk of invasive and metastatic behavior as an alternative rescue mechanism. While taxanes cause similar effects *in vivo* as VDAs, the underlying cellular mechanisms exerted by these drugs are different. Taxane/bevacizumab combination therapy is still used in the treatment of advanced ovarian cancer and breast cancer - although the FDA revoked their approval for breast cancer in 2011 (27, 28). Our results explain the clinical results obtained with the combination of taxanes plus bevacizumab. The drug combination is applied to pre-treated patients with a progressed, metastatic disease and leads to a significantly increased progression free survival – usually accessed by the size changes in established metastases. However, the effect on overall survival is marginal or not even statistically significant(4). An observation, that is in line with a model of initially increased primary treatment efficacy but subsequently enhanced malignancy.

In our murine models combination of taxanes with anti-angiogenic drugs increased the number of CTCs, as a response to acute hypoxic stress. In various cancers CTC levels have been shown to be highly predictive of overall patient survival (29, 30). Importantly, our results indicate that changes in scheduling of the two drugs might circumvent this problem, or alternatively application of TRP channel inhibitors could block the adverse effect by protecting tumor vessels from the vascular disrupting action.

## Acknowledgement

We gratefully acknowledge funding provided by the DFG (Grant Nos. HE3565/2-1 and HE3565/3-1 to E.H.) and the Wilhelm-Sander Foundation (Grant Nos. 2015.001.01 and 2018.080.1 to E.H.). We also want to thank Erna Kleinschroth (Institute of Anatomy, Universität Würzburg) for help and support with histological analysis, and the Core Unit Fluorescence Imaging of the RVZ for support in fluorescence imaging and data analysis.

## References

1. J. Ma, D. J. Waxman, Combination of antiangiogenesis with chemotherapy for more effective cancer treatment. Mol Cancer Ther 7, 3670–3684 (2008).

2. H. Hurwitz et al., Bevacizumab plus irinotecan, fluorouracil, and leucovorin for metastatic colorectal cancer. N Engl J Med 350, 2335–2342 (2004).

3. D. M. O’Malley et al., Addition of bevacizumab to weekly paclitaxel significantly improves progression-free survival in heavily pretreated recurrent epithelial ovarian cancer. Gynecologic oncology 121, 269–272 (2011).

4. K. Miller et al., Paclitaxel plus bevacizumab versus paclitaxel alone for metastatic breast cancer. N Engl J Med 357, 2666–2676 (2007).

5. A. Sandler et al., Paclitaxel-carboplatin alone or with bevacizumab for non-small-cell lung cancer. N Engl J Med 355, 2542–2550 (2006).

6. A. G. Taghian et al., Paclitaxel decreases the interstitial fluid pressure and improves oxygenation in breast cancers in patients treated with neoadjuvant chemotherapy: clinical implications. Journal of clinical oncology: official journal of the American Society of Clinical Oncology 23, 1951–1961 (2005).

7. E. Pasquier et al., Antiangiogenic concentrations of paclitaxel induce an increase in microtubule dynamics in endothelial cells but not in cancer cells. Cancer Res 65, 2433–2440 (2005).

8. D. S. Grant, T. L. Williams, M. Zahaczewsky, A. P. Dicker, Comparison of antiangiogenic activities using paclitaxel (taxol) and docetaxel (taxotere). Int J Cancer 104, 121–129 (2003).

9. G. Bocci, A. Di Paolo, R. Danesi, The pharmacological bases of the antiangiogenic activity of paclitaxel. Angiogenesis 16, 481–492 (2013).

10. J. M. Roodhart et al., Late release of circulating endothelial cells and endothelial progenitor cells after chemotherapy predicts response and survival in cancer patients. Neoplasia 12, 87–94 (2010).

11. F. C. Bidard et al., Clinical value of circulating endothelial cells and circulating tumor cells in metastatic breast cancer patients treated first line with bevacizumab and chemotherapy. Annals of oncology: official journal of the European Society for Medical Oncology / ESMO 21, 1765–1771 (2010).

12. Y. Shaked et al., Rapid chemotherapy-induced acute endothelial progenitor cell mobilization: implications for antiangiogenic drugs as chemosensitizing agents. Cancer Cell 14, 263–273 (2008).

13. A. B. Nair, S. Jacob, A simple practice guide for dose conversion between animals and human. J Basic Clin Pharm 7, 27–31 (2016).

14. S. A. Stewart et al., Lentivirus-delivered stable gene silencing by RNAi in primary cells. Rna 9, 493–501 (2003).

15. A. Klingberg et al., Fully Automated Evaluation of Total Glomerular Number and Capillary Tuft Size in Nephritic Kidneys Using Lightsheet Microscopy. J Am Soc Nephrol 28, 452–459 (2017).

16. R. Szulcek, H. J. Bogaard, G. P. van Nieuw Amerongen, Electric cell-substrate impedance sensing for the quantification of endothelial proliferation, barrier function, and motility. J Vis Exp 10.3791/51300 (2014).

17. G. Carpentier (2018) Angiogenesis Analyzer for ImageJ.

18. G. Carpentier et al., Angiogenesis Analyzer for ImageJ - A comparative morphometric analysis of “Endothelial Tube Formation Assay” and “Fibrin Bead Assay”. Scientific reports 10, 11568 (2020).

19. H. J. de Jonge et al., Evidence based selection of housekeeping genes. PLoS One 2, e898 (2007).

20. L. Silwal-Pandit et al., The Longitudinal Transcriptional Response to Neoadjuvant Chemotherapy with and without Bevacizumab in Breast Cancer. Clin Cancer Res 23, 4662–4670 (2017).

21. V. K. Mootha et al., PGC-1alpha-responsive genes involved in oxidative phosphorylation are coordinately downregulated in human diabetes. Nat Genet 34, 267–273 (2003).

22. A. Subramanian et al., Gene set enrichment analysis: a knowledge-based approach for interpreting genome-wide expression profiles. Proc Natl Acad Sci U S A 102, 15545–15550 (2005).

23. W. C. Liang et al., Cross-species vascular endothelial growth factor (VEGF)- blocking antibodies completely inhibit the growth of human tumor xenografts and measure the contribution of stromal VEGF. J Biol Chem 281, 951–961 (2006).

24. F. Rohrig et al., VEGF-ablation therapy reduces drug delivery and therapeutic response in ECM-dense tumors. Oncogene 36, 1–12 (2017).

25. F. Schutze et al., Inhibition of Lysyl Oxidases Improves Drug Diffusion and Increases Efficacy of Cytotoxic Treatment in 3D Tumor Models. Scientific reports 5, 17576 (2015).

26. Y. Chen et al., Clinical pharmacology of axitinib. Clin Pharmacokinet 52, 713–725 (2013).

27. M. Ratner, FDA panel votes to pull Avastin in breast cancer, again. Nat Biotechnol 29, 676 (2011).

28. S. Hamel et al., Off-label use of cancer therapies in women diagnosed with breast cancer in the United States. Springerplus 4, 209 (2015).

29. Z. B. Xie, L. Yao, C. Jin, D. L. Fu, Circulating tumor cells in pancreatic cancer patients: efficacy in diagnosis and value in prognosis. Discov Med 22, 121–128 (2016).

30. G. Heller et al., The Added Value of Circulating Tumor Cell Enumeration to Standard Markers in Assessing Prognosis in a Metastatic Castration-Resistant Prostate Cancer Population. Clin Cancer Res 23, 1967–1973 (2017).

